# Roughness or sliminess: What makes a haptic material unpleasant

**DOI:** 10.64898/2026.09.05.749471

**Authors:** Müge Cavdan, Zhong Jian Chee, Rochelle Ackerley, Constanze Hesse, Knut Drewing

## Abstract

The current study aims to identify materials that most people experience as being unpleasant to touch and to uncover the perceptual attributes that link to unpleasantness. We sampled 79 everyday materials covering a wide range of sensory properties such as fluidity, granularity, roughness, and softness, e.g., cooking oil, sand, coconut, and silk. We asked participants to freely explore these materials with their dominant hand and subsequently rate their unpleasantness and 23 sensory attributes (e.g., elasticity, stickiness) in the absence of visual and auditory feedback. We collected data on material unpleasantness from two populations: Germany and the UK. Our wide array of materials showed different unpleasantness levels that correlated with specific perceptual material attributes. In general, viscous and deformable materials (e.g., hair gel, play-doh) felt more unpleasant to touch than surface-soft and smooth materials. Results were comparable for the German and UK samples. Overall, our results revealed a relationship between specific sensory attributes of materials and their unpleasantness, which seems to be consistent across German and UK samples. Our findings contribute to the understanding of how various sensory attributes are related to haptic unpleasantness. These insights advance our understanding of haptic perception with various implications, such as product design and affective computing systems, as both seek to integrate touch for detecting and modulating human affective states.

## I. Introduction

TOUCH plays a powerful role in how we judge objects, what they are made of, and how we feel about them. Grasping a metal railing or pressing against sticky, sweaty gym equipment can immediately trigger discomfort or even disgust. These haptic experiences shape our preferences, influencing whether we are drawn to or repelled by something. Touching different materials evokes affective responses that span a spectrum from pleasant to unpleasant (e.g., [1]–[4]) and from painful to innocuous (e.g., [5]–[8]). While pleasant touch and the pain spectrum have been widely studied, unpleasant touch has received little attention. Yet, despite its negative affective impact, unpleasant touch is important as it often provokes strong physiological and behavioral responses. In the present study, we seek to identify materials that are consistently experienced as unpleasant to touch and to uncover the perceptual attributes that contribute to this.

Touch helps us to recognize and identify different objects and materials [9], as well as evoking affective responses when we interact with them [3]. These two aspects of touch are often described in terms of discriminative and affective processing, respectively, although these nearly always occur together [10]. While the former enables the perception of attributes such as texture, temperature, and shape (i.e., sensory aspects), the latter reflects the affective qualities of haptic experiences (i.e., feelings and emotions), like how pleasant or aversive a material feels [11]. Over the last few years, research has increasingly focused on the affective component of touch to uncover what material properties are associated with (un)pleasantness (e.g., [12]–[15]). Findings from this work consistently point to softness (or compliance, the physical correlate) as a key factor: the softer and more compliant a material is, the more pleasant it tends to feel [2], [13]. This effect is especially pronounced when a material mimics the mechanical properties of human skin [12]. However, softness alone does not capture the full spectrum of materials and their properties. Other properties are also related to affective responses. For instance, warmer materials are typically preferred over colder ones [16]; smoother surfaces are rated more positively than rougher ones [8], [17]–[19]; and increased viscosity is associated with decreased pleasantness [20], [21]. These examples illustrate well that discriminative and affective touch are closely intertwined. However, most of these studies have used custom-designed materials that vary only along a single property (e.g., compliance [8]) or a small set of natural materials ([2], [4], [22], [23], see [1] for an extensive work on pleasant materials). While such approaches allow for experimental control and have advanced our understanding of haptic pleasantness, they constrain the ecological validity and generalizability of the findings to unpleasant natural everyday materials.

To address this gap, we devised an experiment in which we used a wide range of everyday materials. While many haptic experiences appear to be universal, such as similar haptic softness perception in Türkiye and Germany [24], [25], recent work suggests that culture can influence how people explore and describe tactile properties [26], [27]. Specifically, individuals from different cultural backgrounds may employ distinct haptic exploratory procedures (i.e., stereotypical hand movements used to explore certain objects [28]) to convey the same tactile attributes [27]. Such findings raise the possibility of cultural variation in affective touch responses.

We, therefore, conducted our experiment in two countries to examine the generalizability and stability of the unpleasantness of materials across independent samples and testing environments (i.e., the UK and Germany). In two separate blocks, participants haptically explored 79 everyday materials (e.g., flour and cling film) and evaluated (a) how unpleasant they felt to touch and (b) how strongly 23 different sensory properties applied to them (e.g., granularity). We then determined the perceptual qualities of the materials with a Principal Component Analysis (PCA) on the adjective ratings. We used a Mann-Whitney *U* test to compare unpleasantness ratings between the UK and Germany, while the correlation analyses of PCA-derived perceptual dimensions and unpleasantness ratings identified which properties drove unpleasantness. This overall approach provides an ecologically valid picture of what people find unpleasant to touch in everyday haptic interactions, and provides insights into how specific perceptual qualities relate to unpleasantness.

## II. Methods

### A. Participants

The sample size was determined based on previous studies that utilized similar exploratory analyses [24], [29]. Consequently, 30 (25 females, *M*_age_ = 24 years, age range: 20-33, *SD*_age_ = 3.49) participants were recruited from Justus Liebig University, Germany, and 30 (20 females, *M*_age_ = 23 years, age range: 19-34, *SD*_age_ = 3.6) from the University of Aberdeen, UK. None of the participants reported any sensory, motor, or cutaneous problems. They were compensated 8 C/£8 per hour or course credit for their time. The study was ethically approved by LEK FB 06 in Giessen (approval 2022-003) and by the Psychology Ethics Committee in Aberdeen (approval code 2212193) following the Declaration of Helsinki [30], excluding the preregistration. Participants provided written informed consent prior to experimentation.

### B. Setup, materials, and adjectives

Noise-canceling headphones (Giessen: Sennheiser HD 4.50 BTNC Aberdeen: Sony WH-1000XM4) were used to mask sounds that could be generated during exploration of the materials. The headphones were also used to present beep sounds that signaled the start and end of a trial, as well as continuous river sounds for masking. A monitor was placed to the left of the participants to present the adjectives and instructions (Fig. 1), and a number keypad was placed next to it to collect responses. A laptop was placed next to the experimenter to run the experiment and collect the responses. The experiment was programmed in Matlab 2022a (MathWorks Inc., 2007) with Psychtoolbox routines [31], [32]. In the Giessen experiment, the hand movements of the participants were recorded with a Sony Digital 4K Video Camera. However, these data will not be presented further in the current study.

**Fig. 1.**
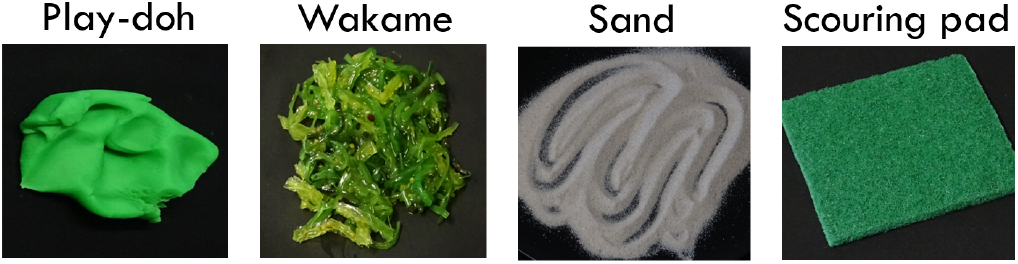
Example stimuli used in the experiment. From left to right: play-doh, wakame, sand, and scouring pad.

**Fig. 2.**
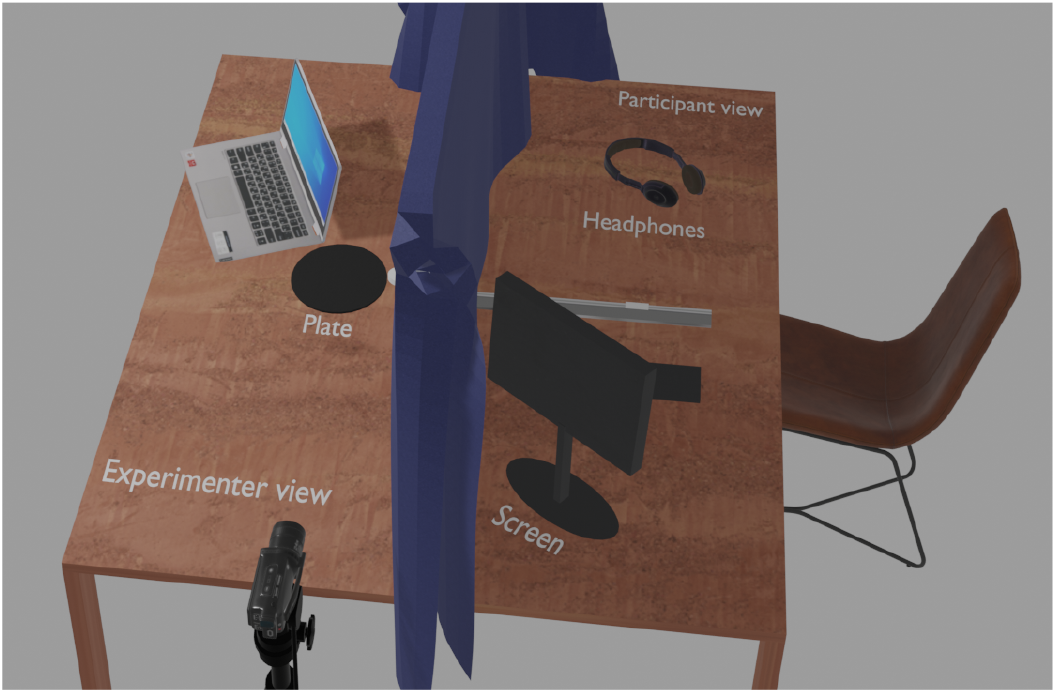
Illustration of the experimental setup in Germany. Please note that this image is rendered in Blender and may not reflect the true sizes of the setup.

During the experiment, participants were seated at a table. A horizontally rotatable armrest was installed on a platform on the table to ensure the materials were explored from a similar distance and to reduce potential arm strain. A curtain was used to block the materials from the participant’s view. The experimenter was seated behind the curtain, and plates (21.5 in diameter, Fig. 1) holding the materials were placed on the table from this side.

In our earlier work (Experiment 1 in [33]), the unpleasantness of a wide range of materials and objects was rated. For the current study, 79 of those were selected, excluding animals and any items that could not be employed due to practical reasons (e.g., raw meat). The stimuli were from everyday items of a wide range of categories, including granular materials (e.g., sand and flour), fluids (e.g., cooking oil and hand cream), and fabrics (e.g., fur and felt; see Supplementary Material or Fig. 3 for the full list). Any materials likely to be substantially altered through exploration were renewed for each participant (e.g., whipped cream and aluminum foil; see Supplementary Material for the full list). The quantity of each material was kept as consistent as possible across participants to avoid differences in how the materials felt (e.g., changes in perceived liquidity [34]).

**Fig. 3.**
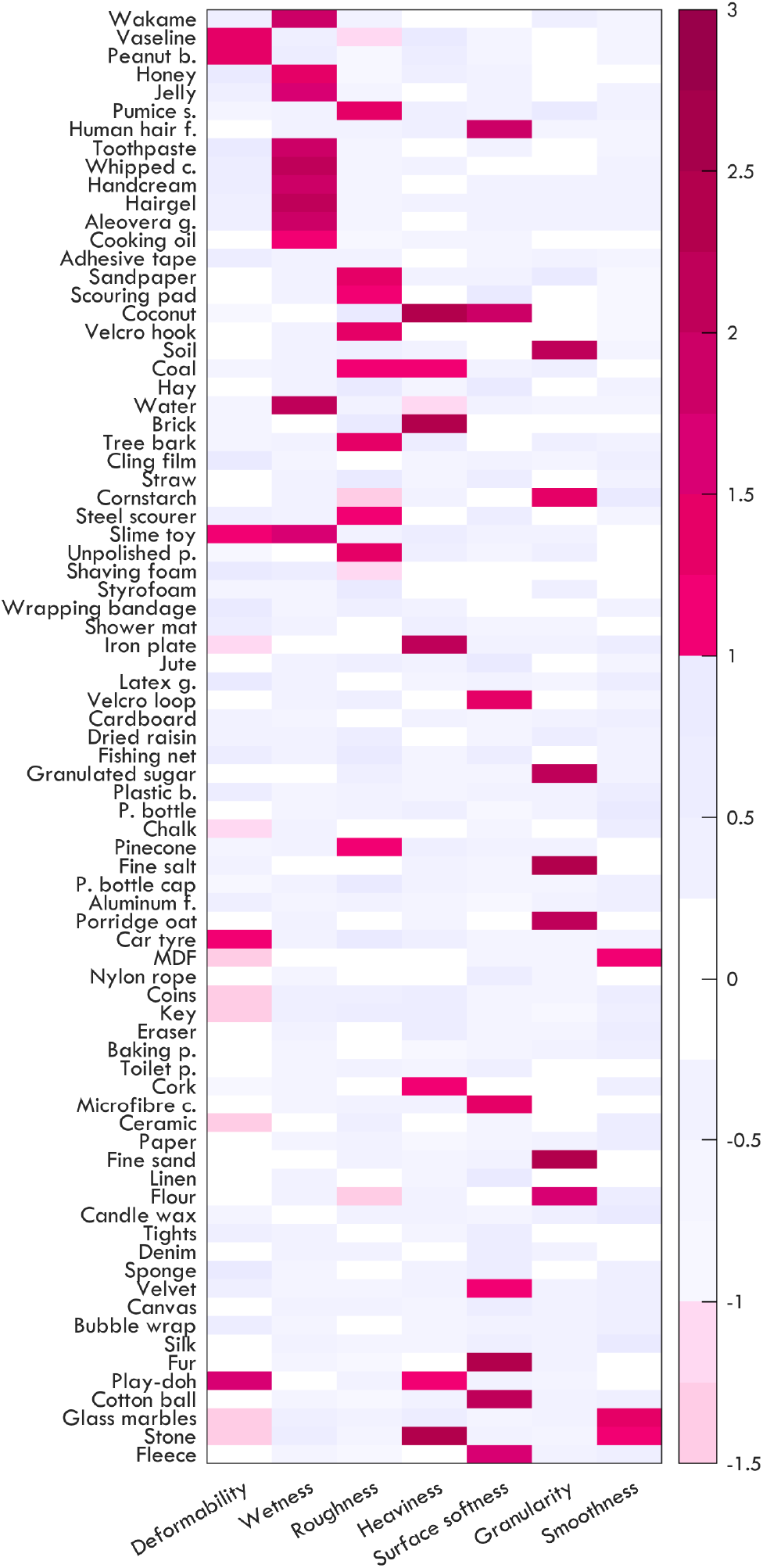
Rotated component scores of materials in each perceptual dimension (i.e., Bartlett scores across Germany and the UK): deformability, wetness, roughness, heaviness, surface softness, granularity, and smoothness; ordered by unpleasantness. Darker, saturated colors indicate positive loadings, and desaturated, lighter colors show negative loadings. Light gray and white areas indicate that loadings were larger than −1 standard deviation or smaller than 1 standard deviation.

We selected 23 sensory adjectives from [35] and [36] covering the dimensions of fluidity (*moist, sticky, slippery*), fibrousness (*hairy, fluffy, fibrous, soft*), roughness (*rough, jagged, smooth*), granularity (*grainy, coarse, powdery*), heaviness (*heavy, light, hard, bulky*), deformability (*elastic, deformable, inflexible*), and temperature (*cold, warm, cool*). The adjectives selected from [35] were already available in both German and English. The remaining two adjectives (i.e., *bulky and cool* from [36]) were translated into German by three German-English bilingual speakers.

### C. Design and procedure

After providing written informed consent, participants performed three practice trials to get familiarized with the setup and the experiment. During the practice trials, they explored and rated the unpleasantness and sensory properties of the practice material (i.e., a woodblock).

This was followed by the main experiment. During the experiment, each participant rated 79 materials according to unpleasantness and 23 sensory adjectives. Specifically, they indicated how unpleasant a material felt (1: *very pleasant*, 7: *very unpleasant*) and how much each sensory adjective applies to a material (1: *not at all*, 7: *very*).

In each trial, once the experimenter had placed a material on the plate, they pressed a button to initiate the trial. This triggered a beep, signaling the start of a 4-second exploration period. During this time, participants explored the material freely, without any movement restrictions. A second beep marked the end of the exploration period and participants were instructed to disengage at that point. Following this, participants first rated how unpleasant the material felt, and then rated the list of adjectives using the number keypad with their left hand. The order of materials and adjective presentations was randomized. Participants could take breaks between trials, and withdraw and clean their hands if needed, especially after exploring certain materials (e.g., peanut butter and cooking oil).

### D. Data analysis

To determine the perceived properties of the materials, we analyzed the individual adjective ratings using covariance-based PCA with varimax rotation, conducted on the combined data from Germany and the UK. Prior to this, we assessed the consistency between adjective ratings using Cronbach’s *α* and evaluated the suitability of the data for PCA using the Kaiser-Meyer-Olkin (KMO) measure and Bartlett’s test of sphericity. Parallel analysis was used to determine the number of dimensions to extract by comparing the empirical eigenvalues to those obtained from randomly generated data of the same size over 1000 iterations. We then computed individual component scores per material using Bartlett method and extracted components separately for each individual. Finally, we averaged these scores across participants for each country, yielding a single value per material and dimension, resulting in 79 (materials) *×*7 (dimensions) data points per country.

We tested the similarity of the dimension structure in the German and the UK populations using Procrustes analysis on these averaged Bartlett scores (see Supplementary Fig. 1 for country-specific Bartlett values). This analysis is a shape comparison technique that assesses the similarity between two configurations by optimally scaling, rotating, and translating one configuration to best match the other. The resulting distance quantifies the dissimilarity, with smaller values indicating greater similarity between the structures [37].

Next, we tested whether the unpleasantness ratings differed across the German and UK samples. To this end, individual unpleasantness ratings were first averaged across participants, yielding one unpleasantness value per material for each country. These values were then submitted to a Mann-Whitney *U* test (as the normality was violated) to check whether unpleasantness ratings varied in our populations.

Since we found evidence for similar unpleasantness across populations, for a coherent presentation of the results, we averaged the unpleasantness scores across German and UK ratings per material (Fig. 3). To determine which materials were perceived as unpleasant, we followed a bootstrapping approach where we approximated a null distribution by shuffling material labels within participants 10,000 times and recalculated mean scores as we did with the observed data. Then, from the 10,000 pseudo means, we extracted the lower and upper critical values for our Omnibus test. The top 5% of lower and upper critical values yields the critical value for our Omnibus test. Therefore, any material with more extreme ratings than these critical values was deemed to be significant (*p <* .05). We additionally computed, for each material, the percentage of participants whose individual ratings exceeded this critical value, as an index of inter-individual agreement (Fig. 4).

**Fig. 4.**
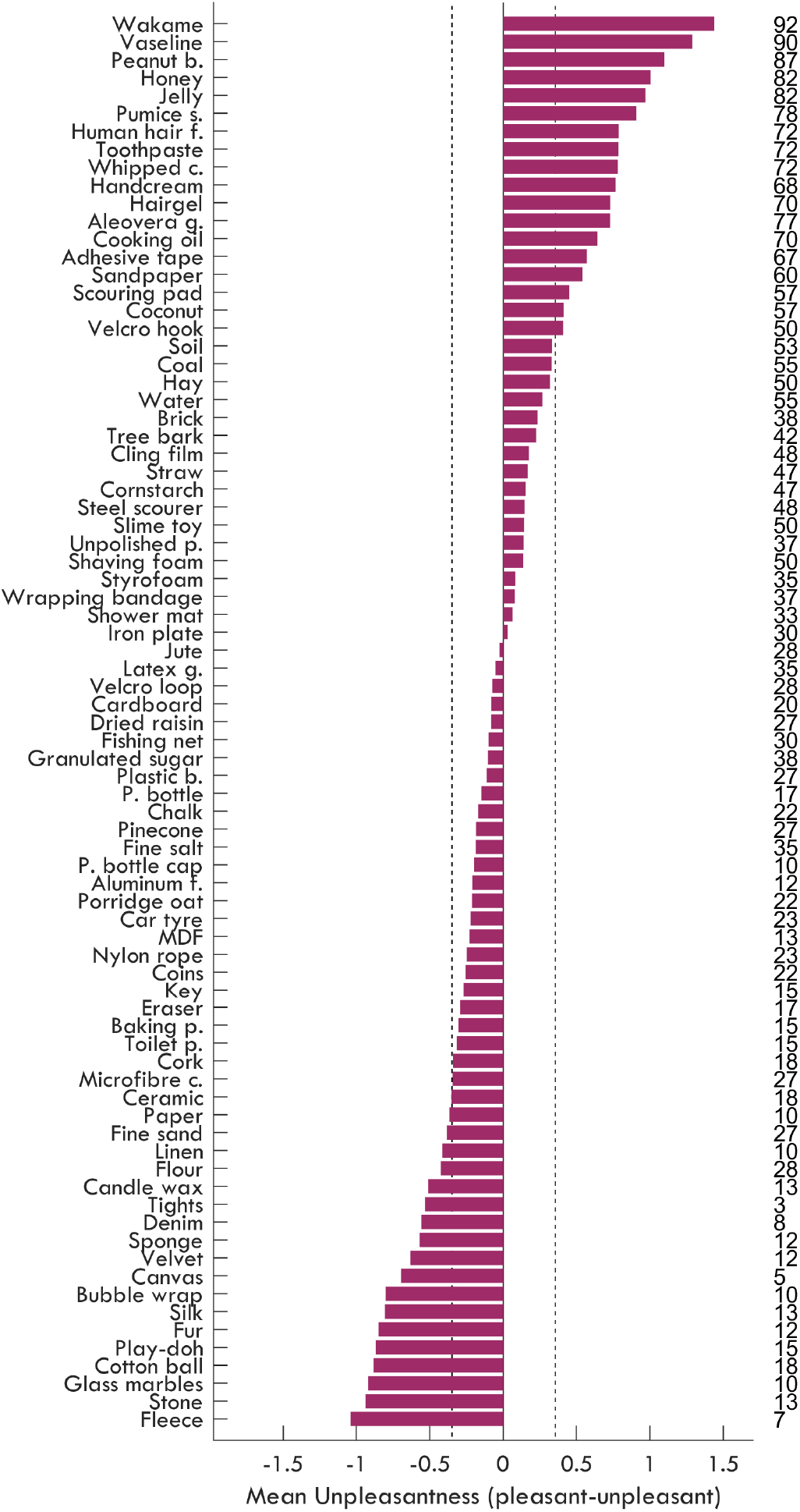
Mean standardized unpleasantness across participants for each material. Negative values indicate pleasantness, while positive values correspond to unpleasantness. Dashed lines indicate the cut-off values. Numbers on the right-hand side of each bar indicate the percentage of participants who rated the material as unpleasant at the individual level (i.e., *z >* .355).

In order to investigate which material properties relate to unpleasantness, we correlated the mean Bartlett values for materials identified as unpleasant or pleasant (obtained from Germany and the UK) in bootstrapping analysis with the mean unpleasantness ratings for each dimension (i.e., 37 materials *×*7 dimensions, Fig. 5). As we had specific hypotheses (see Results C) for each dimension and correlation, we did not correct the p-values.

**Fig. 5.**
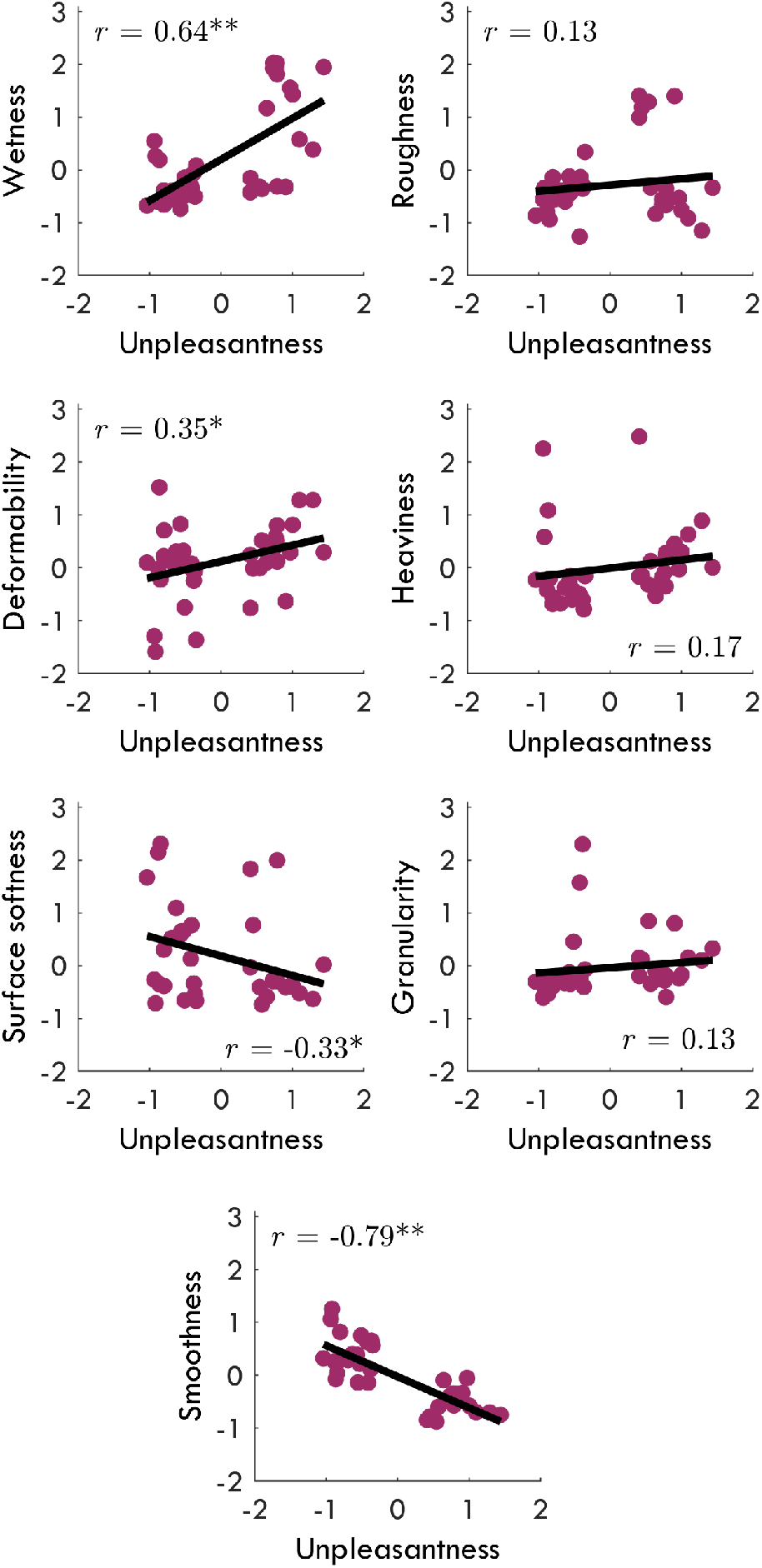
The relationship between perceptual dimensions and unpleasantness — x-axes indicate mean unpleasantness, while y-axes values show mean Bartlett scores of unpleasant and pleasant materials obtained from Germany and the UK: wetness, roughness, deformability, heaviness, surface softness, granularity, and smoothness, respectively. Please note that only unpleasant and pleasant materials are plotted following the correlations.^∗∗^*p <* .001, ^∗^*p <* .05.

The PCA was conducted in SPSS (Version 28.0), while the Mann-Whitney *U* test was performed in JASP (Version 0.16.2.0). The Procrustes, bootstrapping, and correlation analyses were performed in MATLAB using custom scripts (R2022b).

## III. Results

### A. Perceptual dimensions obtained from exploring affective materials

First, we determined the structure of the material space obtained from Germany and the UK. To begin, we assessed the consistency of individual ratings for 79 materials across 23 adjectives, in order to determine the extent to which individuals responded similarly to the materials. Cronbach’s *α* revealed excellent consistency between participants in both the German (*α* = .864 - .996) and the UK (*α* = .778 - .994) samples [38], supporting the inclusion of all participants’ data in subsequent analyses.

Next, we determined the dimensionality of perceived materials by submitting individual ratings from Germany and the UK to a covariance-based PCA. KMO score of .834 indicated good sampling adequacy, and Bartlett’s test of sphericity (*χ*^2^(253) = 47394.153, *p <* .001) revealed that the observed correlations between perceptual ratings were meaningful.

**TABLE I.** Rotated factor loading of adjectives from the PCA.

| Adjective | Deformability | Wetness | Roughness | Heaviness | Surface Softness | Granularity | Smoothness |
| --- | --- | --- | --- | --- | --- | --- | --- |
| deformable | **1.914** | 0.190 | -0.276 | -0.501 | 0.112 | -0.034 | 0.109 |
| elastic | **1.418** | 0.053 | 0.187 | -0.030 | 0.087 | -0.494 | 0.405 |
| inflexible | *-1.388* | -0.111 | 0.808 | 0.959 | -0.302 | -0.053 | 0.500 |
| hard | *-1.129* | -0.172 | 1.032 | 1.065 | -0.437 | 0.014 | 0.472 |
| soft | **0.965** | 0.253 | -0.880 | -0.412 | 0.909 | 0.058 | 0.693 |
| cool | -0.163 | **1.626** | 0.032 | 0.024 | -0.051 | -0.141 | 0.192 |
| cold | -0.129 | **1.568** | 0.039 | 0.148 | -0.090 | -0.154 | 0.066 |
| moist | 0.528 | **1.530** | -0.565 | -0.014 | -0.230 | -0.083 | -0.307 |
| slippery | 0.329 | **1.226** | -0.584 | -0.014 | -0.349 | 0.015 | 0.646 |
| sticky | 0.784 | **1.034** | -0.542 | 0.146 | -0.317 | 0.066 | -0.354 |
| rough | -0.179 | -0.433 | **1.538** | 0.246 | 0.195 | 0.631 | -0.402 |
| jagged | -0.146 | -0.211 | **1.407** | 0.239 | 0.092 | 0.183 | -0.083 |
| heavy | -0.249 | 0.138 | 0.165 | **1.283** | -0.019 | -0.107 | 0.111 |
| light | 0.262 | -0.148 | 0.165 | *-1.661* | 0.174 | 0.106 | 0.705 |
| bulky | -0.093 | -0.016 | 0.313 | **1.486** | 0.048 | 0.044 | 0.237 |
| fibrous | 0.036 | -0.235 | 0.565 | -0.148 | **1.513** | 0.259 | -0.199 |
| hairy | -0.031 | -0.194 | 0.119 | 0.081 | **1.138** | -0.092 | -0.191 |
| fluffy | 0.286 | -0.081 | -0.400 | -0.165 | **1.246** | 0.097 | 0.369 |
| grainy | -0.138 | -0.099 | 0.547 | -0.056 | 0.008 | **1.597** | -0.161 |
| powdery | -0.042 | -0.072 | -0.138 | -0.141 | 0.075 | **1.000** | 0.137 |
| coarse | -0.213 | -0.118 | 0.861 | 0.171 | 0.125 | **1.283** | -0.216 |
| smooth | 0.028 | 0.620 | -0.885 | 0.091 | -0.314 | -0.342 | **1.435** |
| warm | 0.317 | -0.249 | 0.038 | 0.077 | 0.385 | 0.197 | 0.444 |

**Note:** Grayed-out adjectives correspond to the highly loading adjectives per dimension: positive loadings are highlighted in bold while negative loadings are italicized.

The parallel analysis identified seven components; accordingly, we extracted seven components, explaining 70.7% of the total variance. The first rotated component, which we termed *deformability*, accounted for 13.2% variance with adjective loadings *deformable, elastic, inflexible, hard, and soft*. The second rotated component explained 12.9% of the variance with the adjectives of *cool, cold, moist, slippery, and sticky*, which we termed *wetness*. We named the third rotated component *roughness* with the adjective loadings *rough and jagged*, which accounted for 12.2% of the variance. The fourth rotated component explained 11% of the variance with adjectives *heavy, light, and bulky*, which we called *heaviness*. The fifth rotated component explained 8.1% of the variance with the adjective loadings *fibrous, hairy, and fluffy*, which we called *surface softness*. The sixth rotated component was *granularity* with the adjectives *grainy, powdery, and coarse*, which explained 7.3% of the variance. Finally, the seventh rotated component was *smoothness*, which explained variance of 6.0% and *smooth* loadings (see Supplementary Fig. 1).

We next assessed the degree of structural similarity between Germany and the UK by performing a Procrustes analysis on Bartlett scores of materials across all dimensions. Based on this analysis, we calculated the residual sum of squared errors after mapping the configurations between Germany and the UK. The error between countries was low (.10), which shows a good fit ([39], [40]) between the derived material dimensions in Germany and the UK. To formally test significance, we followed a bootstrapping approach [41]. We generated 10,000 pseudo Bartlett values (Supplementary Fig. 1) by shuffling the empirical Bartlett scores separately for each country. We then calculated the Procrustes errors for each material and pseudo-comparison. The empirical mapping error between Germany and the UK falls within the lowest 2.5th percentile of the pseudo error distribution, indicating a high degree of structural similarity between countries.

### B. Material unpleasantness in German and the UK populations

We conducted a Mann-Whitney *U* test to compare unpleasantness ratings between the German and UK populations. The analysis revealed no significant difference in haptic unpleasantness between the two groups (*U* = 3649.00, *p* = .07). This suggests that there was no meaningful difference between Germany and the UK in the haptic unpleasantness of materials.

We z-standardized and collapsed the pleasantness data obtained from Germany and the UK and identified the unpleasant and pleasant materials as those falling above (.355) or below (-.349) the cutoff values, respectively, as determined by the bootstrapping method (see Supplementary Fig. 2 for country-specific means). Unpleasant materials were *wakame, vaseline, peanut butter, honey, jelly, pumice stone, human hair (fake), toothpaste, whipped cream, hand cream, hair gel, aloe vera gel, cooking oil, adhesive tape, sandpaper, scouring pad, coconut*, and *velcro hook side*, while the pleasant ones were *ceramic pieces, paper, fine sand, linen, flour, candle wax, tights, denim, sponge, velvet, canvas, bubble wrap, silk, fur, play-doh, cotton balls, glass marbles, stone (polished)*, and *fleece* (see Fig. 5 for standardized and Supplementary Fig. 2 for raw data).

### C. The relationship between perceptual qualities and affective responses

We tested the relationship between perceptual dimensions and unpleasantness, expecting wetness [21], roughness [8], [17], [18], and heaviness to correlate positively, and deformability, surface softness, granularity [4], and smoothness to correlate negatively.

Expectedly, unpleasantness was positively related to wetness (*r* = .64, *p <* .001) and negatively to surface softness (*r* = -.33, *p* = .048) and smoothness (*r* = -.79, *p <* .001). Unexpectedly, deformability was positively correlated with unpleasantness (*r* = .35, *p* = .034) while the other relationships were not statistically significant (roughness: *r* = .13, heaviness: *r* = .17, granularity: *r* = .13, all *p >* .05).

## IV. Discussion

We investigated how unpleasant everyday materials are felt in German and UK populations and which material properties are associated with unpleasantness. Moving beyond earlier studies that relied on custom-made samples varying along a single material dimension or focused mainly on pleasant touch, we systematically sampled a broad range of naturally occurring materials spanning a wide spectrum of (un)pleasantness, thus establishing a richer and more ecologically valid basis for studying unpleasant touch. Both populations exhibited comparable unpleasantness ratings for the materials tested, with items such as Vaseline, sandpaper, and pumice stone consistently evaluated as unpleasant. The materials varied along seven dimensions: deformability, wetness, roughness, heaviness, surface softness, granularity, and smoothness. Consistent with previous work, unpleasantness increased with increasing wetness [16], [20], [21], [42] and decreasing surface softness [4] and smoothness [8]. Interestingly, in contrast to earlier findings [4], [12], greater deformability was also associated with increased unpleasantness. These observed quality-unpleasantness relations are relevant for affective computing, where understanding how touch elicits affect is critical for designing effective haptic and interaction systems.

### A. The relationship between perceptual qualities and unpleasantness

The correlation analyses revealed that materials perceived as higher in wetness and as more deformable tended to feel unpleasant to touch, whereas surfaces that were soft and smooth were generally experienced as pleasant. This is also reflected in the set of materials rated as the most unpleasant: *algae, vaseline, peanut butter, honey, jelly, pumice stone, human hair (fake), toothpaste, whipped cream, hand cream, hair gel, aloe vera gel, cooking oil, adhesive tape, sandpaper, scouring pad, coconut*, and *velcro hook side*. These findings largely align with previous work showing that soft and smooth materials are typically associated with pleasant touch, while viscous materials are not [1]–[3], [7], [17], [18], [21]. However, our results challenge prior conclusions relating to softness. While earlier studies reported that softness was consistently linked to pleasant touch, the deformable materials in our study were often judged as unpleasant. Several factors may explain this discrepancy. First, previous research has usually represented softness with a single class of solid stimuli (e.g., foam, sponge, silicone). By contrast, in daily life, we encounter non-solid materials that are also “soft” or deformable, such as hair gel or sand [24], [25] ([43] for a review), yet do not necessarily evoke the same hedonic response. Second, materials rarely vary along a single perceptual dimension. For example, toothpaste can be described not only as viscous, but also as deformable and cold, and it is likely the combination of these properties that shapes its unpleasantness. When we examine Bartlett’s results (Fig. 3), it becomes apparent that many of the materials that loaded in the deformability dimension in our study deform under load in ways characteristic of substances lying between liquidity [44] and viscoelasticity [45]. This suggests that softness cannot be treated as a universally pleasant quality, but rather depends on how it is expressed across multiple material properties. In other words, the (un)pleasantness of touch emerges not from softness alone, but from the specific constellation of features - such as viscosity, temperature, and deformation. Therefore, while considering the unpleasantness of everyday materials, it is important to consider their multidimensional nature and draw conclusions accordingly.

### B. Cross-country consistency in unpleasantness

Interestingly, perceptual and unpleasantness judgments were strikingly similar across these populations and individuals. This cross-cultural agreement suggests that the mapping between certain material properties (e.g., viscosity, roughness) and their affective evaluation is not strongly shaped by cultural learning, but may instead reflect more fundamental aspects of haptic processing. In other words, the unpleasantness associated with viscosity or roughness, or the pleasantness of certain materials/objects, may arise from shared sensory and affective mechanisms rather than being shaped by culture. Such consistency implies that unpleasantness judgments are not highly idiosyncratic but generalize across individuals, which is crucial for applications in product design or affective computing.

Systems that rely on haptic interactions can therefore exploit these stable associations to predict user responses with rather little need for individual adaptation. Nevertheless, it is important to note that our sample included only two Western European populations, and future work should extend these findings to a broader set of cultural contexts.

### C. Relevance of unpleasantness

Understanding unpleasantness itself is important as negative haptic experiences strongly influence how people interact with materials and products. Whereas pleasant touch has often been emphasized in the literature (e.g., [11]), unpleasantness may be equally influential, shaping avoidance behaviors [46], [47] or emotional responses [48] in daily life. Importantly, aversive haptic experiences include subtle yet powerful sensations (e.g., non-painful but uncomfortable temperatures), which can evoke strong affective reactions. While these responses might have evolutionary value in signaling potential threats or contamination [49], they also extend to everyday contexts, where they influence consumer choices, interpersonal interactions, and clinical conditions characterized by tactile hypersensitivity. This highlights the importance of considering not only the rewarding aspects of touch but also its aversive nature, which can drive material rejection, inform ergonomic considerations, and guide therapeutic practices. The present paper contributes to this effort by providing materials that can be used in psychological studies and affective computing to systematically examine the unpleasant aspects of haptic experiences.

While our focus was on affective responses to inanimate materials, the neural processes supporting these evaluations likely overlap with those involved in social touch. Xu et al. demonstrated that mechanoreceptive A primary afferents can discriminate fine-grained characteristics of naturalistic social touch at behaviorally relevant timescales. These same afferents encode haptic information such as pressure, deformation, and temporal dynamics. Thus, the mechanoreceptive pathways that allow the nervous system to distinguish subtle variations in stroking velocity or contact force during social interactions would also encode material properties such as viscosity, deformability, and texture. This shared low-level encoding provides the sensory foundation for affective evaluations of both social and material/object-based touch. Our findings, therefore, complement work on social affective touch by demonstrating that specific physical attributes of non-social materials systematically shape unpleasantness.

In the current study, we provide an overview of materials that individuals consistently rated as unpleasant to touch (Fig. 4, unpleasantness agreement at the individual level reflected on percentages). However, it is important to note that such judgments are stimulus-set dependent. Similarly, the unpleasantness of a material is relative to the stimulus set. Therefore, the same material might be pleasant within a stimulus set while unpleasant within another one. For instance, materials that might be expected to feel unpleasant, such as tree bark [4], were not rated as strongly unpleasant, water - typically assumed to be more neutral or even pleasant [21]-was judged more towards the unpleasant side when evaluated within a more diverse sample. Taken together, our results provide a nuanced picture of relative unpleasantness across a diverse set of materials.

Future work should extend these findings by testing more diverse cultural groups, including different body sites, and by combining behavioral measures with physiological or neural indices of affective touch. Such work will further clarify how perceptual dimensions of materials are transformed into affective responses and how these signals can be integrated into technologies that sense, predict, and modulate human touch experiences.

## V. Conclusion

The present study identified which everyday materials are consistently experienced as unpleasant to touch and revealed the perceptual dimensions that underlie these evaluations. Across German and UK participants, we found a stable seven-dimensional structure of material perception, and comparable unpleasantness ratings, indicating that the affective evaluation of common materials generalizes across these populations. Unpleasantness was most strongly linked to higher viscosity and deformability, as well as reduced surface softness and smoothness, showing that “soft” materials are not uniformly pleasant and that unpleasant touch emerges from specific combinations of haptic attributes. These insights provide an ecologically valid overview of unpleasant haptic materials and offer a useful basis for applications in product design and affective computing, where predicting and modulating unpleasant touch experiences is critical.

## Acknowledgments

The authors would like to thank Kimberly Glas for her assistance with data collection, setup, and material organization. They also thank Aleksandra Mijailovic and Tim-Luca Schmitz for their help with data collection in Giessen, as well as Daniela Ruseva for their support with data collection in Aberdeen.

## VI. Biography Section

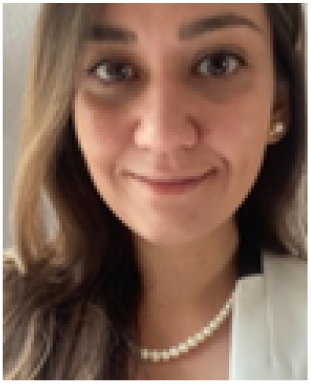

**Mü ge Cavdan** is a postdoctoral researcher at Justus Liebig University Giessen since 2021. She earned her Ph.D. in experimental psychology from the same university, focusing on visual and haptic material perception. She was awarded the World Haptics Best Student Paper Award in 2019 and the Best Paper in EuroHaptics 2020 & 2024. In 2022 & 2024, she received the Innovation in Haptics supported by the IEEE Technical Committee on Haptics of the IEEE Robotics and Automation Society. Her research interests include material perception, active and affective touch, timing, and multisensory processing.

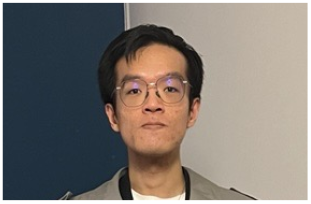

**Zhong Jian Chee** is a postdoctoral researcher at the University of Aberdeen since 2023. He earned his Ph.D. in psychology and cognitive neuroscience from the University of Nottingham Malaysia, focusing on autistic traits, executive functions, and musical sophistication. His current research interests include material perception and affective touch, particularly how unpleasant-to-touch materials influence human actions and behaviors.

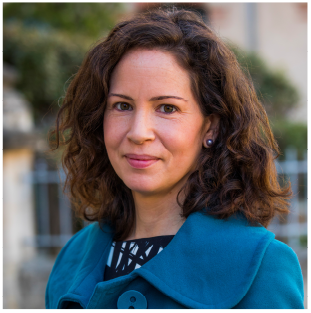

**Rochelle Ackerley** obtained her Ph.D. in physiology from the University of Bristol in 2006, then worked as a cognitive neuroscientist in industry, before completing postdoctoral work at the University of Manchester, UK on sensorimotor control and the University of Gothenburg, Sweden on touch. She joined the CNRS in 2017, and was promoted to Research Director in 2023, where she is based at Aix-Marseille University. Her research examines how touch shapes behavior in humans, using both neurophysiological and psychophysical approaches, including focuses on affective touch and temperature.

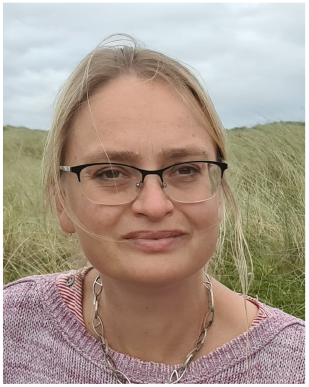

**Constanze Hesse** received her PhD in experimental psychology from Justus Liebig University Giessen, Germany. She subsequently held postdoctoral research positions at Ludwig-Maximilians-Universität Munich and at Durham University (UK). In 2012, she joined the University of Aberdeen (Scotland) as a Lecturer, where she was promoted to Senior Lecturer in 2017 and to Professor in 2022. Her research examines how visual and cognitive factors shape motor behavior, and how the physical and affective properties of materials influence intentions to act.

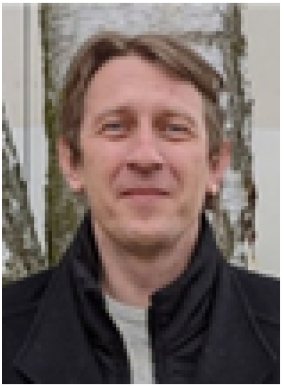

**Knut Drewing** received the Ph.D. degree in psychology from Ludwig Maximilian University of Munich, Germany, in 2001, and the postdoctoral lecture qualification (habilitation) in psychology from JLU Giessen, Germany, in 2010. He worked in the MPIs for Psychological Research (Munich) and for Biological Cybernetics (Tuebingen). He is an Associate Editor-in-Chief for IEEE Transactions on Haptics. He is currently a Professor at the HapLab, Department of Psychology, JLU Giessen. His research interests include haptic perception, multisensory integration, and time perception.

