## Supplemental file for "Roughness or sliminess: What makes a haptic material unpleasant"

**Materials.**We used the following materials and objects in the experiment: iron plate, nylon rope, sandpaper, toilet paper*, creamy peanut butter*, whipped cream*, aluminum foil*, baking paper*, porridge oats, fishing net, aloe vera gel*, latex wipe/gloves, tights, Play-doh, plastic bag, microfiber cloth, jelly*, plastic bottle caps, plastic water bottle, glass marbles, dried raisins, candle wax, pumice stone, cork, unpolished pebbles, eraser, stone, cooking oil, pine cone, tree bark, slime toy, soil, hay (dried grass), fake hair, wrapping bandage, bubble wrap*, linen, denim, silk, anti-slip shower mat, fur, fleece, canvas, cardboard, MDF (medium density fiberboard), cling film*, jute, wakame* (kept in the fridge), hair gel*, sponge, coins, key, granulated sugar, A4 paper*, fine salt, hand cream, fine sand, water, scouring pad, adhesive tape (gluey side)*, chalk, coal, steel scourer, Velcro hook side, Velcro loop side, styrofoam, car tyre, toothpaste, straw (plant), Vaseline*, ceramic, cotton ball, velvet, shaving foam*, honey*, brick, cornstarch, coconut, and flour. Asterisks indicate that the stimulus was replaced for each participant.

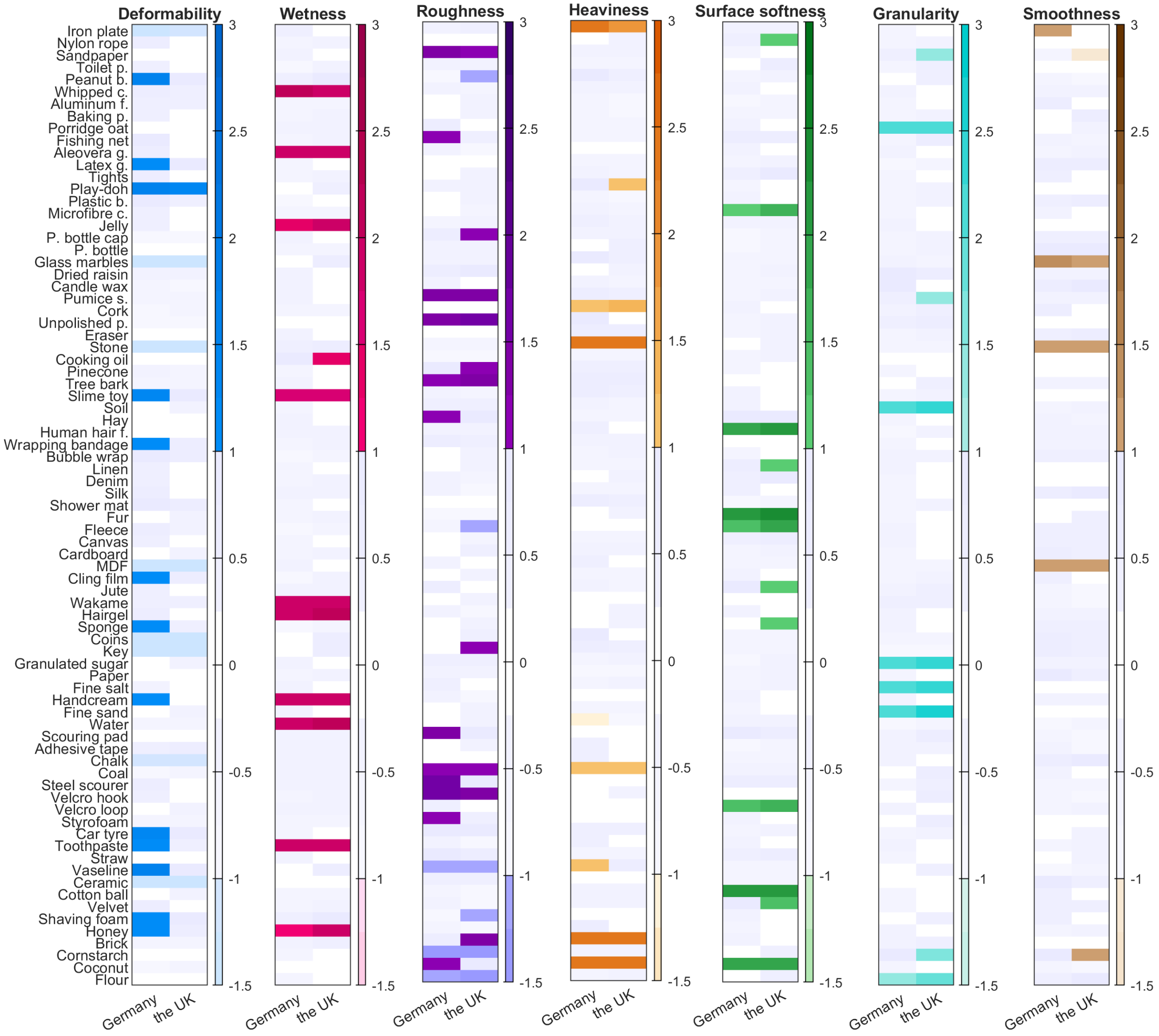


Figure 1: Supplementary Figure S1. Rotated component scores of materials in each perceptual dimension: deformability, viscosity/temperature, roughness, heaviness, fibrousness, granularity, and smoothness. Darker, saturated colors indicate positive loadings, and desaturated, lighter colors show negative loadings. Light gray and white areas indicate that loadings were larger than -1 standard deviation or smaller than 1 standard deviation.

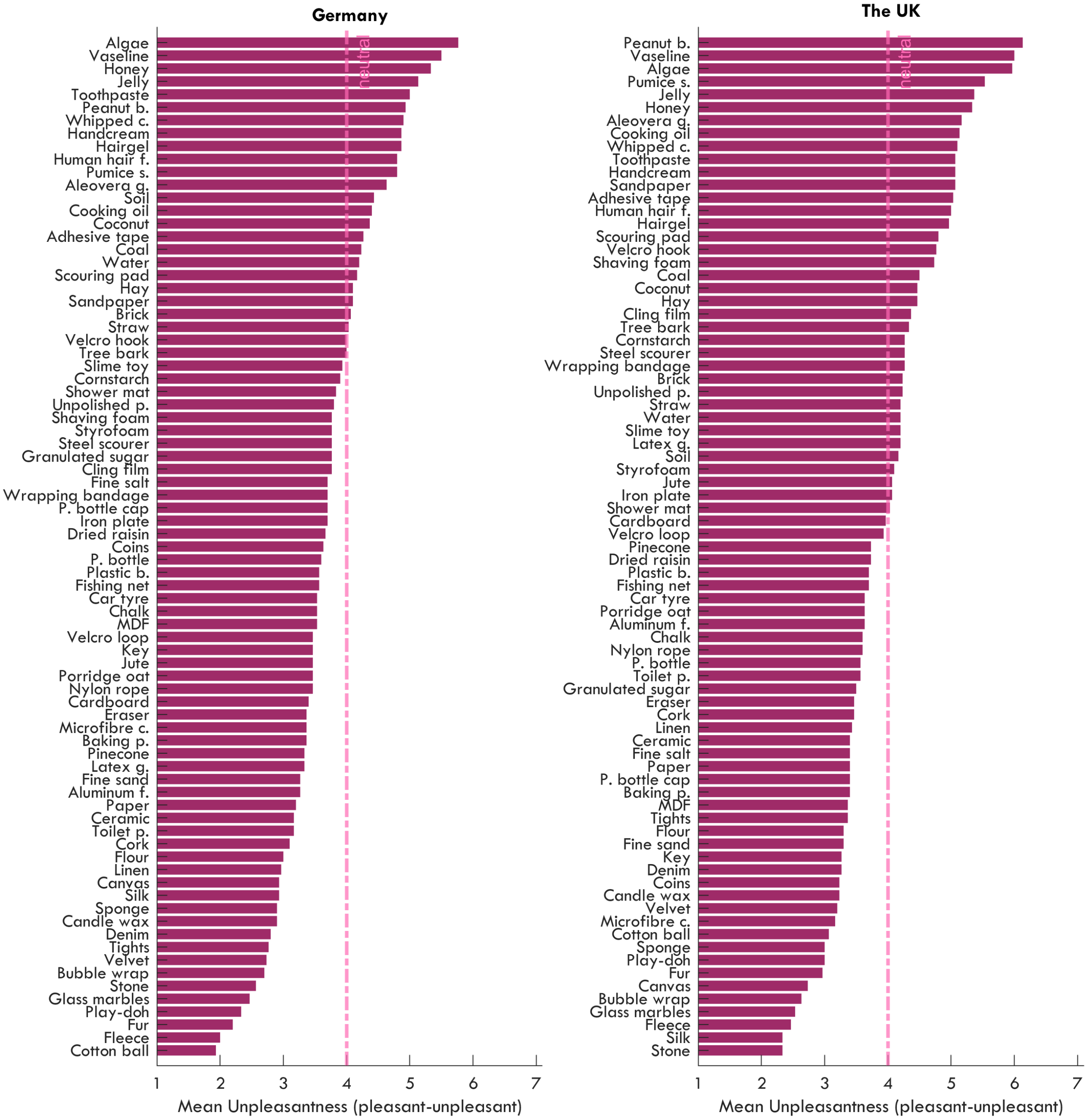


Figure 2: Supplementary Figure S2. Raw unpleasantness scores across materials for Germany and the UK. Dashed lines indicate the neutral point.
